# FlexENN: A Graph Neural Network for Binding Energy Prediction of Globular and Intrinsically Disordered Proteins

**DOI:** 10.64898/2026.05.15.725440

**Authors:** Maryum Irshad, Kassandra Ori-Mckenney, Ruxandra I. Dima

**Affiliations:** Department of Chemistry, University of Cincinnati, 45221 Cincinnati, Ohio, United States; Department of Molecular and Cellular Biology, University of California, Davis, California, United States

## Abstract

Intrinsically disordered proteins (IDPs) drive a large fraction of cellular signaling, transcription and assembly through interfaces that lack a single defined geometry, making the prediction of their binding energies beyond the reach of methods calibrated on the rigid complementarity of folded complexes. Here we introduce an adaptive message passing graph neural network, FlexENN, that predicts binding energies across the full structural spectrum, from folded domains to complexes with one disordered partner. The FlexENN architecture constructs a local interface graph in which each residue node integrates relative geometry, dynamic descriptors, and sequence information, allowing the network to infer conformational fuzziness from a single structure. Benchmarked across folded complexes and IDP systems, including tubulin-tubulin interfaces within microtubules (MTs) and MTs in complex with microtubule associated proteins (MAPs), with reference binding affinity values extracted from our experiments, the model retrieves physically meaningful energetic profiles for complexes where rigid-body methods fail. More broadly, this work delivers an accurate, structure-aware approach to predicting binding free energies for the broad class of disordered-mediated complexes formed by IDPs.

## Introduction

Intrinsically disordered proteins (IDPs) and intrinsically disordered regions (IDRs) represent a major class of biomolecules that defy the classical structure-function paradigm. ^1^ Rather than adopting a single stable three-dimensional structure, IDPs exist as dynamic ensembles of rapidly interchanging conformations, enabling them to sample a wide range of structural states.^2,3^ This intrinsic flexibility facilitates their central roles across a broad spectrum of complex cellular processes, including transcriptional regulation, ^4,5^ cellular signaling, ^6^ and genome maintenance,^7,8^ where adaptability and responsiveness to environmental cues are essential. However, the same conformational freedom that drives the IDP functions also contributes to their involvement in neurodegenerative diseases, where the lack of a stable structural framework increases susceptibility to misfolding and pathological aggregation. ^9^

The functions of IDPs are not governed by a fixed structure but by a sequence-ensemble-function relationship, in which amino acid composition and patterning encode a distribution of conformations with distinct physicochemical properties. ^10,11^ Features such as charge distribution, hydrophobicity, aromatic interactions, and transient secondary structure collectively determine ensemble properties including chain compaction, interaction accessibility, and multivalency.^12–14^ These properties enable IDPs to engage in diverse binding modes, ranging from partial folding upon interaction to highly dynamic fuzzy complexes and fully disordered bound states. Consequently, IDP-mediated interfaces are often diffuse, heterogeneous, and spatially extended, with binding driven by transient, distributed contacts rather than well-defined structural complementarity. ^15,16^

This ensemble-driven nature of the IDP interactions introduces a fundamental challenge for modeling binding energetics. In contrast to folded proteins, where a single conformation can often approximate the binding interface, IDP interactions emerge from a distribution of conformations and context-dependent interactions, including sensitivity to post-translational modifications, solvent conditions, and partner identity. As a result, the binding affinity is dependent on a combination of local physicochemical environments, residue-level flexibility, and multivalent interaction patterns.

In contrast to the dynamic and ensemble-driven nature of the IDP interactions, existing computational methods for predicting protein-protein binding affinities have largely been developed for folded proteins. Early approaches such as DFIRE^17^ relied on knowledge-based statistical potentials derived from atomic contact frequencies observed in experimentally characterized protein structures, estimating the binding energetics through comparisons to an idealized reference state. This geometric focus was further refined by models such as PRODIGY,^18^ which demonstrated that relatively simple features, including interfacial contacts and non-interacting surface areas, can effectively approximate binding affinity, reinforcing the central role of geometry.

Subsequent developments in the field have increasingly incorporated machine learning and deep learning techniques to capture more complex interaction patterns. RoseTTAFold^19^ introduced a three-track neural network that integrates 1D sequence information, 2D residue-residue distance maps, and 3D coordinate-based representations, enabling simultaneous reasoning across multiple structural scales. Beyond protein-protein interactions, RoseTTAFold has been widely applied to tasks such as de novo protein structure prediction and design,^19–21^ mutation effect analysis,^22^ and foldability assessment,^23^ underscoring its versatility in integrating multi-scale representations within computational structural biology. In parallel, AtomNet^24^ incorporated atomic-level evolutionary information alongside structural potentials to achieve high-resolution interface scoring, though at the cost of substantial computational expense and a reliance on rigid-body assumptions.

More recent efforts have focused on enhancing the feature representations and leveraging multimodal learning strategies. PerSpect-EL^25^ utilized persistent spectral theory and Hodge Laplacian-based descriptors to encode topological and geometric characteristics of protein interfaces, while Yang et al.^26^ extended the contact-based features used by PRODIGY^18^ with physicochemical surface properties, demonstrating improved performance through ensemble modeling.

Deep learning architectures have further advanced the field by integrating learned sequence representations with structural graph-based models. Methods such as ProAffinity-GNN,^27^ PPI-Graphomer, ^28^ and SSIF-Affinity^29^ combine protein language model embeddings with graph neural networks to capture both local interfacial interactions and border contextual information. Nevertheless, these frameworks rely on comprehensive representations of entire protein complexes and construct extensive graphs that include many regions, especially those distal to the binding interface, under the assumption that proteins exist in folded, structurally well-defined states.

Another related class of approaches focuses on leveraging biophysical descriptors derived from empirical energy functions, particularly those computed using Rosetta. Ferraz et al. ^30^ employed a range of machine learning models, including support vector regression, XGBoost, and an artificial neural network, trained on Rosetta-derived terms such as van der Waals interactions, solvation energies, and electrostatics to predict absolute binding free energies. Similarly, the PBEE^31^ framework utilized a super learner ensemble that integrates predictions from multiple machine learning algorithms, achieving improved accuracy through the combination of diverse predictive models. This approach relies on a fixed set of 52 Rosetta-based features computed across a set of folded protein complexes. While these methods demonstrated strong performance, their reliance on predefined energy terms and folded protein structures limits their applicability to IDP systems.

Despite all these advances, most existing approaches remain focused on folded, structurally well-defined protein complexes, hindering their ability to generalize to IDP systems. This limitation highlights the need for computational frameworks capable of accounting for the structural heterogeneity and conformational flexibility, motivating the development of models specifically tailored to binding affinity predictions of IDPs.

To overcome the limitations of existing methods, we present a novel, inherently adaptive message-passing graph neural network architecture, FlexENN, that accurately predicts the binding energies of both folded and intrinsically disordered proteins. Rather than applying rigid deterministic constraints that heavily penalize coordinate fluctuations, FlexENN conceptualizes the binding interface as a dynamic microenvironment. It achieves this by restricting the graph construction to the interface and integrating relative distance vectors with explicit dynamic structural descriptors, such as B-factors and secondary structure propensities. This framework empowers the network to infer structural fuzziness directly from static representations, contextually weighing flexibility against static geometry on a residue-by-residue basis. Consequently, FlexENN dynamically prioritizes its feature reliance on the inherent structural nature of the complex, leveraging backbone geometry to evaluate structured globular domains, while prioritizing extended spatial proximity to capture the transient contacts characteristic of disordered assemblies. To validate this framework, we benchmarked the model against a diverse set of both folded proteins and IDPs, such as tubulin and microtubule-associated proteins, demonstrating its capacity to extract physically meaningful energetic profiles across a structural spectrum ranging from rigidly folded domains to highly flexible IDPs.

## Materials and Methods

### Datasets Preparation

To ensure sufficient representation of intrinsically disordered proteins, two separate datasets were made: one containing IDPs and another containing folded proteins. In the combined dataset, IDPs constitute only ∼19% of the total samples, resulting in a significant imbalance relative to folded proteins. Such an imbalance can bias machine learning models toward feature characteristics of folded complexes, potentially limiting their ability to capture the distinct interaction patterns of IDPs. Given that the primary goal of this work is to develop a predictive model tailored for IDPs, separating the datasets enables more effective training and evaluation on IDP-specific interactions.

A total of 2798 folded protein complexes were downloaded from PDBbind+ database (v2020.R1),^32^ each with a 3D structure and an experimental binding affinity. The PDBbind dataset is widely used for developing machine learning models for predicting biomolecular binding affinities. However, previous studies have reported several data quality concerns within the protein-ligand dataset from PDBbind, including redundant or duplicate complexes with identical affinity values. These entries can introduce bias during model training and evaluation, particularly when the same structures appear in the training and test sets. In such cases, machine learning models may effectively memorize these repeated samples rather than learning generalized structure-affinity relationships. Consequently, when PDBbind is applied to protein-protein interaction studies, similar redundancy issues may arise. To ensure robust model evaluation and prevent memorization of repeated samples, the dataset was refined by removing (1) duplicate entries, (2) complexes without a valid affinity value, (3) structures with more than 10 chains, (4) pairs of complexes that were structurally identical (with different PDB IDs) and had the same binding affinity. (5) IDPs were also removed from this dataset to make sure it is a set dedicated to folded protein structures. After filtering, 2,257 unique folded protein-protein complexes remained in the dataset (PBEE + Folded).

For the IDPs dataset, we expanded the PBEE dataset,^31^ which originally had 532 protein-protein complexes. Additional IDP complexes were obtained from the DisProt ^33^ database, and their experimental binding affinities were obtained from PDBbind+. To ensure a broad coverage of disorderliness, 703 structures with disordered regions ranging from 5% to 80% were selected. The dataset was filtered using the same criteria applied to the folded proteins, resulting in a final dataset (PBEE + IDPs) of 1,071 IDP complexes.

The experimental affinity for all these complexes was obtained from PDBbind+, and the binding free energy was calculated from the dissociation constant, *K*_*d*_, using the relation, ΔG = −*RT* ln *K*_*d*_, where *R* is the ideal gas constant, and *T* is the temperature in Kelvin (298.15 K was used for all cases). The final datasets used in this study include PBEE, PBEE + Folded, and PBEE + IDPs, which together provide an expanded and structurally diverse set of protein-protein interactions for model training and evaluation. The distribution of binding energy values across these datasets is shown in Figure 1, highlighting that the expanded datasets retain the energetic characteristics of the original PBEE distribution while increasing the coverage.

**Figure 1:**
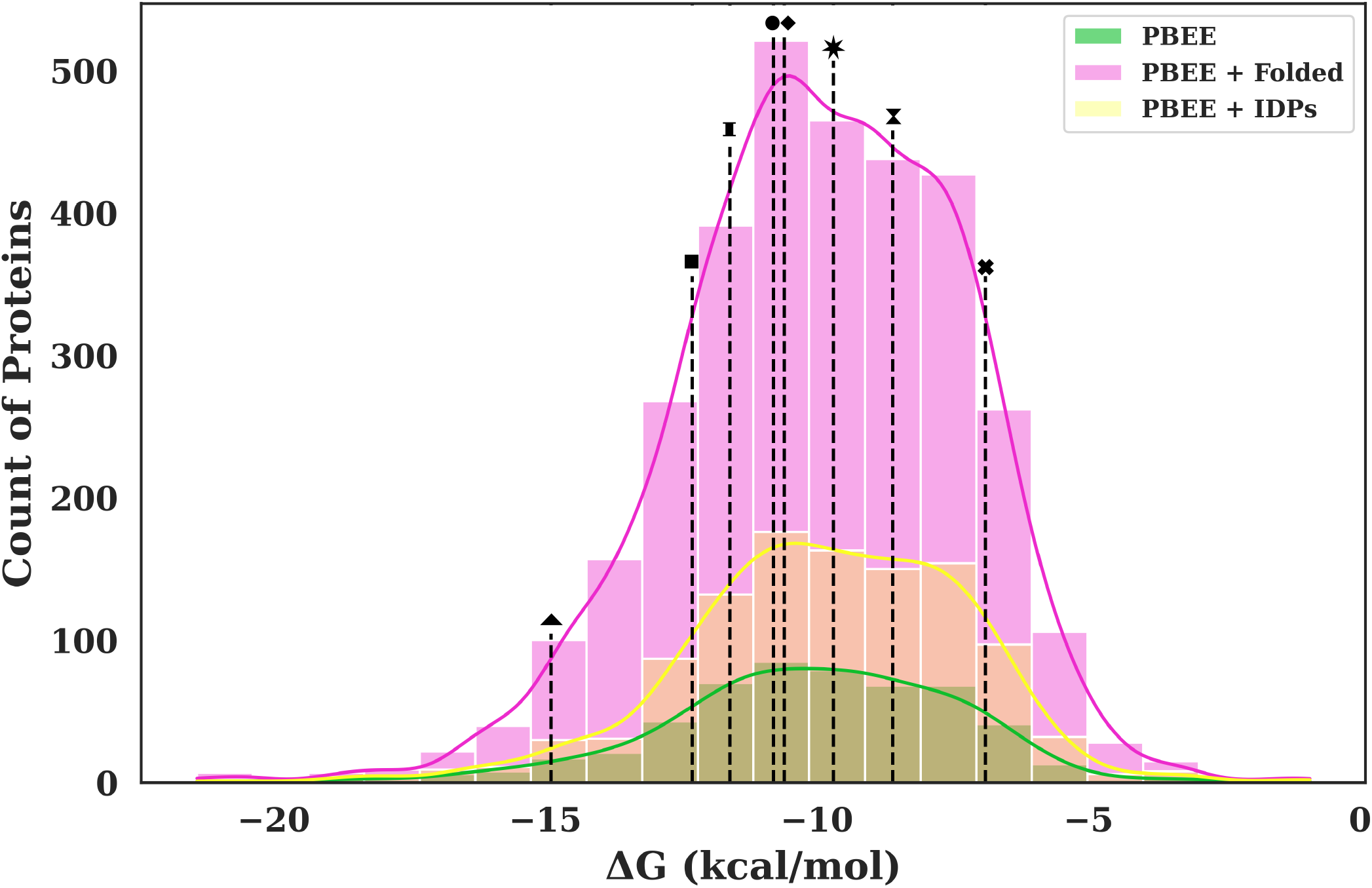
Distribution of binding free energy (ΔG) values across the three datasets used in this study: PBEE (green), PBEE + Folded (pink), and PBEE + IDPs (yellow). All three datasets span a comparable ΔG range (approximately − 20 to − 2 kcal/mol) and are centered around − 10 kcal/mol, indicating that the folded- and IDP-augmented sets preserve the overall energetic character of the original PBEE distribution while increasing the sample size. Dashed vertical lines indicate ΔG values for the benchmark complexes used in this work: Tubulin interdimer interface (▲, −14.9 ± 1.5 kcal/mol), MAP7-MT (■, −12.3 kcal/mol), *β*-catenin-Tcf4 (**I**, −11.6 kcal/mol), DCX-MT (NDC + CDC) (•, −10.8 kcal/mol), Tau-MT (♦, −10.6 kcal/mol), PCNA-p21 (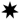, −9.7 kcal/mol), *β*-catenin-Tcf4/BCL9 (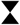, −8.6 kcal/mol), Tubulin lateral interface (×, −6.9 ± 0.4 kcal/mol).

### Benchmark Set

Based on the known redundancy and potential data leakage issues in PDBbind,^34^ we chose to evaluate our models on a benchmark set independent of the training set. The benchmark comprises two complementary subsets to test distinct aspects of model generalization. The first focuses on microtubule (MT) and microtubule-associated proteins (MAPs) interaction landscape, a biologically critical system characterized by a mix of rigid structural interfaces (at the MT tubulin interdimer and lateral interfaces) and intrinsically disordered proteins (IDPs). The second extends the assessment to three additional IDP-partner complexes drawn from distinct biological contexts, chosen to span a range of interface architectures and binding affinities.

For the MT-MAPs subset, we selected four PDB entries representing diverse binding modes: As shown in Figure 2, **6RF2** is a Cryo-EM structure of the C-terminal DC repeat (CDC) of human doublecortin (DCX) bound to MT.^35^ This structure also served as a reference for investigating the MT lattice itself, providing three distinct interaction interfaces for analysis: the strong MT longitudinal interdimer interface, the weaker lateral interfaces of the MT lattice, and the specific CDC-MT binding interface. A model’s ability to correctly rank the energies of MT interdimer and lateral interfaces in the complex was used as a primary criterion for physical plausibility. **6REV**, the Cryo-EM structure of the N-terminal DC (NDC) of human doublecortin (DCX) bound to MT,^35^ complements 6RF2; including both CDC and NDC allows us to assess the models’ capacity to distinguish domain-specific interaction landscape. **6CVN** and **7SGS**, shown in Figure 3, Cryo-EM structures of tau ^36^ and MAP7^37^ bound to MT, respectively, represent interactions involving IDPs. These IDPs present a particular challenge for traditional energetic models that often penalize conformational heterogeneity.

**Figure 2:**
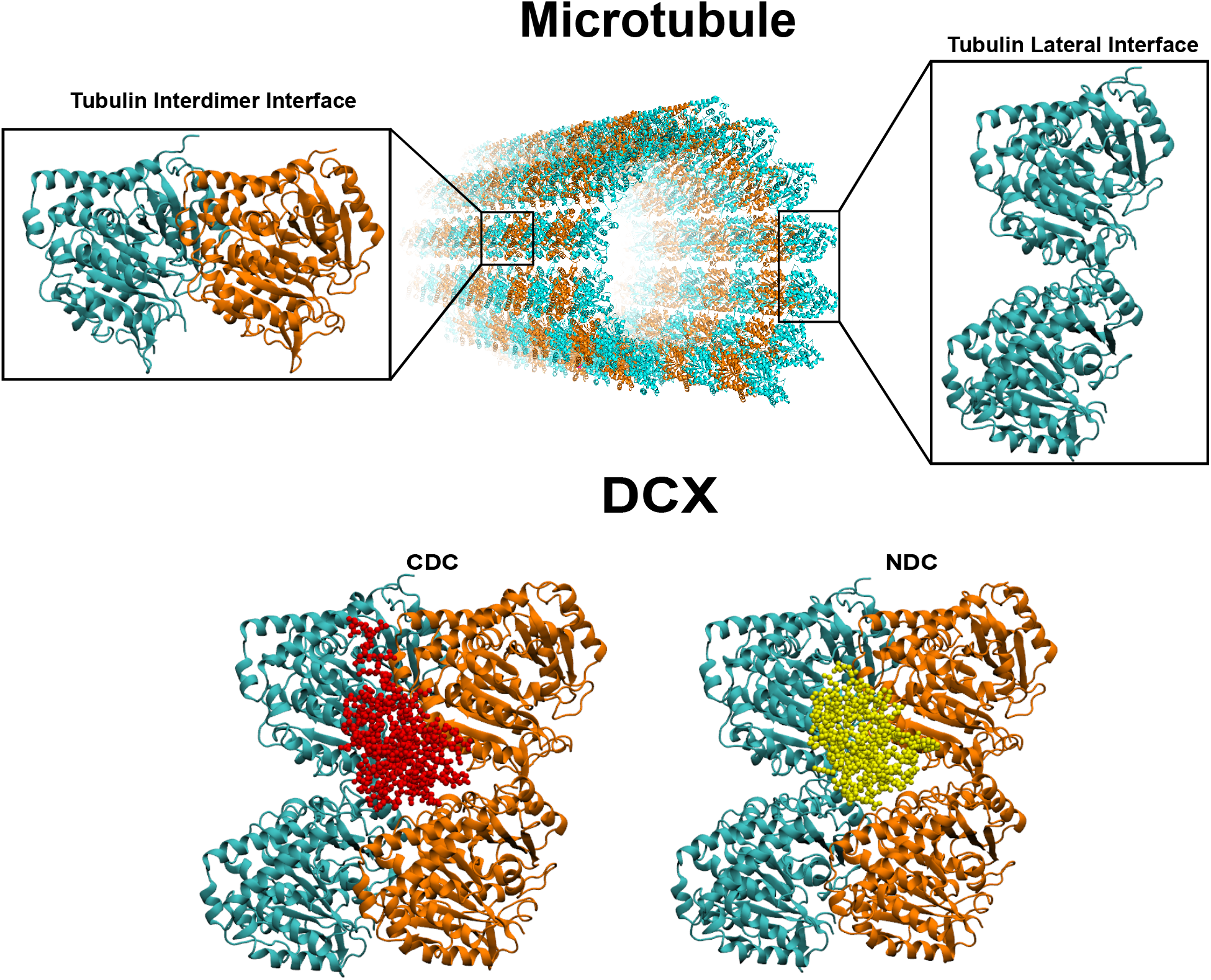
PDB structures of the benchmark set of the MT-MAPs folded proteins.

**Figure 3:**
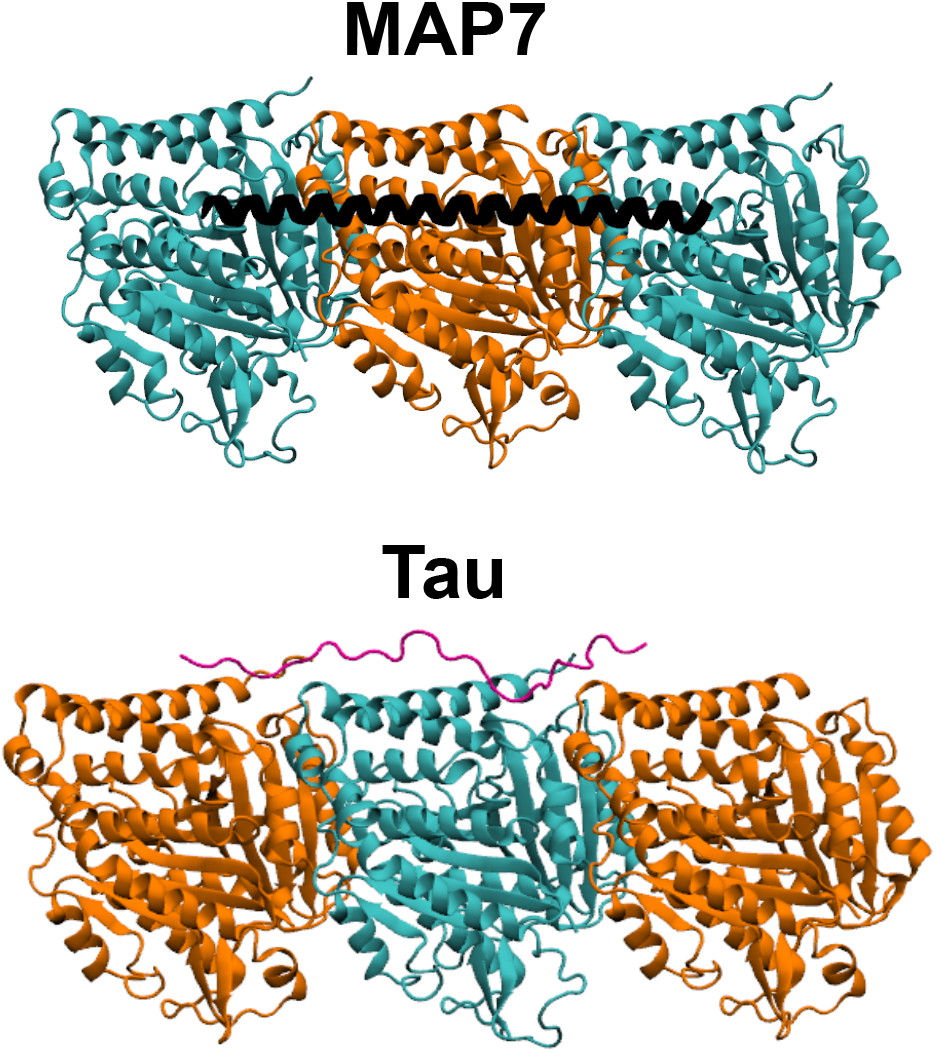
PDB structures of the benchmark set of the MT-MAPs intrinsically disordered proteins.

The folded protein dataset was used to obtain predictions for MT interdimer, lateral, NDC-MT, and CDC-MT interactions. In contrast, the IDPs dataset was used to obtain values for the MAP7-MT and tau-MT setups. We evaluated the models on the experimentally resolved cryo-EM structures of the MAPs obtained from the RCSB Protein Data Bank. ^38^

Collectively, MTs and MAPs play a central role in cytoskeletal dynamics, neuronal development, and intracellular transport. All three MAPs considered in this work, MAP7, tau, and DCX, form envelopes along the MT lattice, shaping the local interaction landscape. ^39^ The concept of a “MAP code” suggests that these MAPs selectively modulate motor protein behavior; for example, MAP7 enhances kinesin-1 motility, while tau and DCX inhibit kinesin-1, thereby biasing transport along MTs.^40,41^ This functional diversity, driven by MAP-MT interactions, makes these systems a compelling and biologically relevant test set for evaluating the performance of binding energy models.

The binding free energies (ΔG) for DCX, MAP7, and tau were derived from equilibrium dissociation constants (*K*_*d*_) determined experimentally using two complementary approaches. First, MT co-sedimentation assays measured binding as a function of microtubule concentration, with bound and unbound fractions quantified by SDS-PAGE and fit to a one-site binding model. Second, total internal reflection fluorescence microscopy (TIRF-M) was used to measure fluorescence intensity of labeled MAPs on immobilized microtubules, with KD obtained by fitting saturation curves to a one-site binding hyperbola. Where cooperative binding was observed (for tau and DCX), Hill coefficients were additionally calculated using Hill-Langmuir fits or analysis of cooperative assembly formation.

The second subset, shown in Figure S1, comprises three IDP-partner complexes that bind to their partners through distinct interface architectures from those in the MT-MAPs subset. **1AXC** is the crystal structure of the human proliferating cell nuclear antigen (PCNA) homotrimer in complex with the C-terminal region of the cyclin-dependent kinase inhibitor, p21.^42^ PCNA forms a sliding clamp that tethers DNA polymerases *δ* and *ϵ* to DNA during synthesis, enabling processive replication, and also functions in nucleotide excision repair. ^43,44^ p21 directly inhibits DNA replication by binding tightly to PCNA via its C-terminal region, thereby preventing polymerase recruitment to the clamp. ^45^ Structurally, the C-terminal region of p21 adopts an extended conformation with a 310-helical turn, anchoring three hydrophobic residues into the inter-domain connecting loop pocket of PCNA. As a localized interaction involving a disordered region with clear biological consequences for cell cycle control, 1AXC provides a stringent test of the model’s ability to capture binding energetics of the short linear motif (SLiM)-mediated interactions.

Both **1JDH** and **2GL7** involve *β*-catenin (Figure S1), the central transcriptional effector of canonical Wnt signaling pathways, bound to disordered partners involved in Wnt-driven gene activation. **1JDH** is the crystal structure of the human *β*-catenin armadillo repeat domain bound to the N-terminal region of the transcription factor Tcf4. ^46^ Tcf4 is an IDP in isolation and folds upon binding to *β*-catenin, being anchored by a salt bridge between Tcf4 glutamate residue and Lys 312 of *β*-catenin. The *β*-catenin-Tcf4 complex is central to the canonical Wnt signaling pathway and is a key driver of transcriptional activity in colon cancer, where Tcf4 is the predominant Tcf family member. ^47^ **2GL7** is the structure of a heterotrimeric complex comprising the human *β*-catenin armadillo repeat, the homology domain 2 (HD2) of BCL9, and the Tcf4 *β*-catenin binding domain.^48^ BCL9 is a transcriptional coactivator essential for *β*-catenin mediated activation of Wnt target genes, and the *β*–catenin–BCL9 interface represents a promising drug target for blocking oncogenic Wnt signaling. ^49^ In this complex 2GL7, Tcf4 and BCL9 do not make direct contact with each other and bind to *β*-catenin independently. Together, 1JDH and 2GL7 test the model’s ability to predict binding energetics for IDP-protein complexes within a shared signaling context but with distinct interface geometries, a binary *β*-catenin-Tcf4 complex versus a heterotrimeric *β*-catenin-BCL9-Tcf4 assembly.

The reference binding energy values for the tubulin interdimer and lateral interfaces were taken from our previous work.^50^ Experimental binding energy values were obtained from the literature for PCNA-p21,^51^ *β*-catenin-Tcf4,^52^ and *β*-catenin-BCL9-Tcf4. ^48^ Together, these complexes in the benchmark span the major classes of IDP-partner interactions, from short motif-mediated binding to extended IDR-scaffold engagement and multi-protein assemblies, providing a stringent test of model generalization across structural diversity that characterizes the disordered proteome.

### Message Passing Geometric Graph Model

Because IDPs lack a stable folded globular structure, traditional structure-based methods fail to predict binding affinities for such systems. To capture the extreme conformational heterogeneity and complex binding thermodynamics of IDPs, we developed a translation-invariant message passing graph neural network, FlexENN. By combining relative spatial geometries with a specialized, biophysically motivated node feature space, the model learns residue-level interaction patterns that contribute to binding energy prediction. A schematic overview of the model architecture is shown in Figure 4.

**Figure 4:**
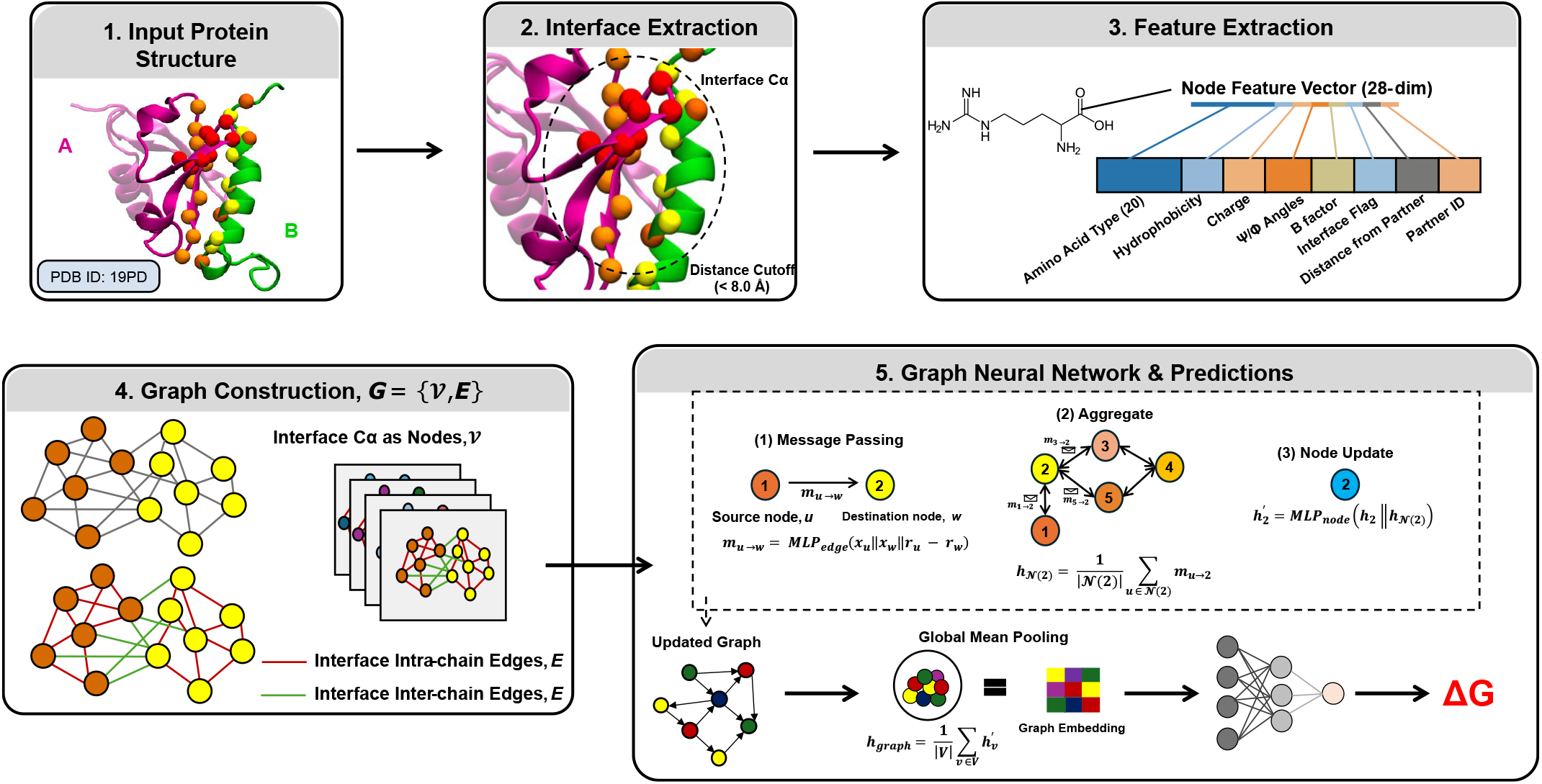
Overview of the FlexENN: the graph neural network architecture used to predict protein-protein binding energy (ΔG) for IDPs. **(1) Input protein.** A protein-protein complex (shown here with PDBID: 19PD; chains A and B shown in magenta and green, respectively). **(2) Interface Extraction**. Interface residues are identified as those with a C*α* atom within an 8 Å distance on partner chains. **(3) Feature Extraction**. Each interface residue is encoded as a 28-dimensional node feature vector comprising amino acid identity (one-hot, 20 dims), hydrophobicity, charge, backbone *ψ*/*ϕ* angles, B-factor, an interface flag, distance to the nearest residue on the partner chain, and partner chain identifier. **(4) Graph Construction**. A graph G = {𝒱, E} is built with interface C*α* atoms as nodes (𝒱); edges (E) connect C*α* of interface residues within the distance cutoff and are labeled as either intra-chain (red) or inter-chain (green). **(5) Graph Neural Network and Prediction**. Node embeddings are refined through successive message-passing layers. At each layer, every edge produces a message computed by an edge-level MLP that takes as input the feature vectors of the source and destination nodes together with their relative coordinates. The messages arriving at each node are then averaged over its neighborhood, and a node-level MLP combines this aggregated neighborhood signal with the node’s current embedding to produce its updated representation. After the final message-passing layer, the resulting per-node embeddings are combined by global mean pooling into a single graph-level embedding, which is passed through a multilayer perceptron to yield the predicted binding free energy ΔG.

### Graph Construction

We represent the protein complexes as residue-level directed graphs ***G* = {𝒱, *E*}**, where ***𝒱*** denotes the set of nodes and **E** is the set of edges. Each node, *v*_*i*_ ∈ ***𝒱*** corresponds to a single amino acid residue, and an edge represents a spatial relationship between Carbon-alpha (C*α*) atoms of two residues. Rather than including the entire protein, the graph is restricted to the binding interface, the region where two protein chains make contact, to focus only on the residues most relevant to binding. Therefore, a residue is included in the graph if its C*α* atom is within a distance of 8 Å from any residue of the interacting partner chain. For example, if a leucine on chain A is located within 8 Å of a glutamate on chain B, both residues satisfy the criterion and are included as nodes in the graph. Residues that are farther from the interface are excluded to reduce computational complexity. Each node *v*_*i*_ is associated with a physicochemical feature vector *x*_*i*_ ∈ ℝ^28^, encoding its chemical and structural properties, and a spatial coordinate vector *p*_*i*_ ∈ ℝ^3^ corresponding to its C*α* position.

The edge set **E**, integrates two distinct types of edges *r* ∈ {1, 2}, defined based on distances between C*α* atoms of the interface residues. The intra-chain edges capture the local geometry and folding constraints within a single chain. They are established between residues *i* and *j* if they belong to the same chain and satisfy ∥*p*_*i*_ − *p*_*j*_∥ < 8 Å. For instance, two neighboring residues along a helix or loop that are spatially close (e.g., 6 Å apart) will be connected. Inter-chain edges capture direct spatial contacts between residues in opposing chains, formed between residues within a cutoff ∥*p*_*i*_ − *p*_*j*_∥ < 4.5 Å. For example, a residue in chain A that sits 4.2 Å from a residue in chain B forms an inter-chain edge, indicating a likely interaction.

Together, these two edge types encode both the backbone structural context within each chain and the spatial proximity between chains at the binding interface, providing the model with a multi-scale geometric representation of the complex. Figure 5 illustrates this graph construction for a representative benchmark system (MAP7-MT, PDB:7SGS), showing both the residue-level contact network and its mapping onto the 3D structure.

**Figure 5:**
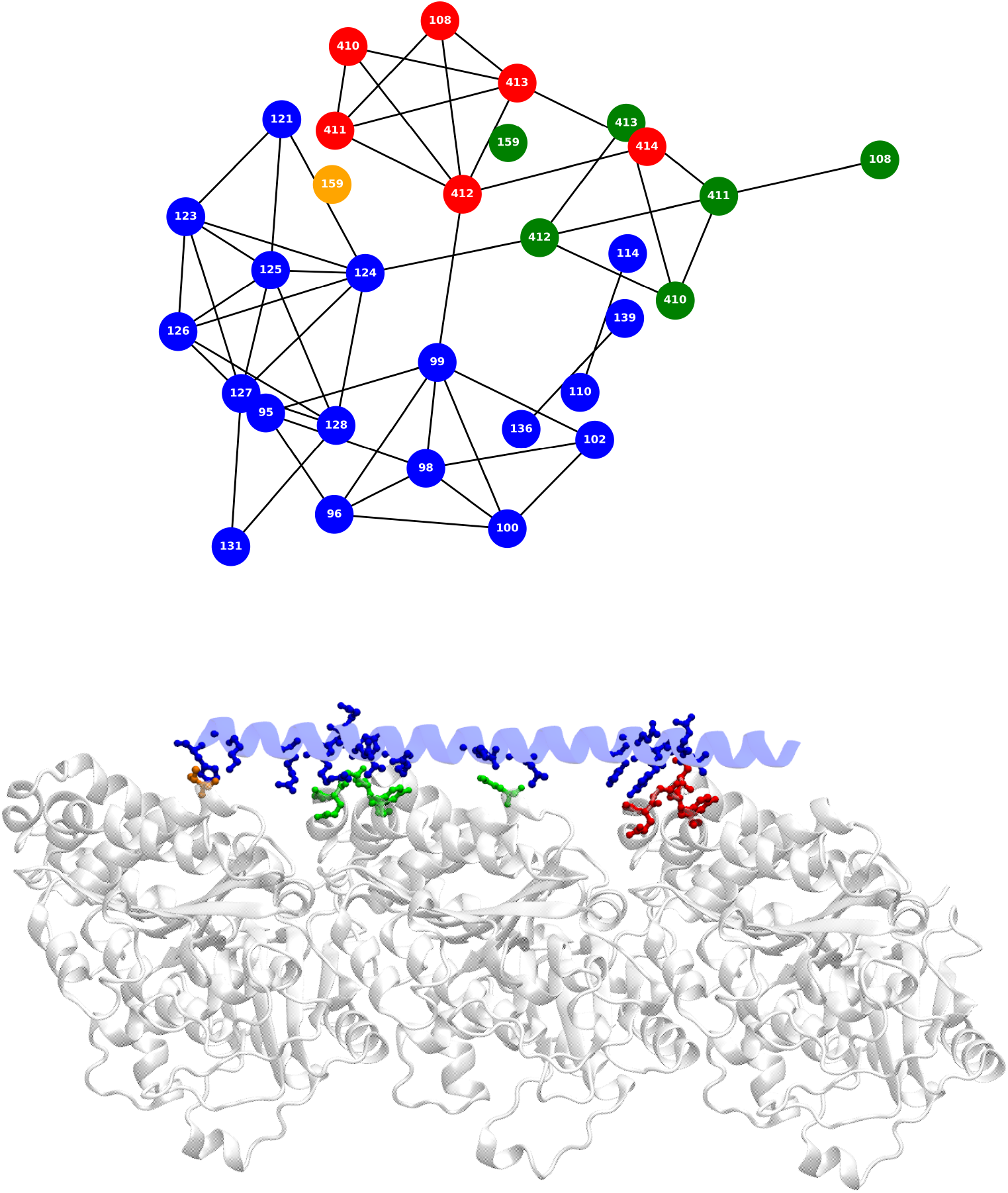
Residue contact network from the FlexENN-I for MAP7 (PDB:7SGS). Top - Interaction graph from FlexENN-I representing the connectivity of residues at the interface. Nodes represent individual amino acid residues, and edges were defined based on spatial proximity using a dual cut-off approach: interchain edges for C*α* of residues within 4.5 Å distance between the partner chains and intrachain edges within 8 Å for the same chains. Node colors denote specific chains: blue for MAP7 (chain A), while orange, green, and red represent MT alpha and beta tubulins. Bottom - structural mapping of the network residues onto the 3D structure. MAP7 (blue helix) is showing interactions across the longitudinal interface of MT dimers (gray cartoon). The residues identified in the graph are shown in stick representation and color-coded by chains similar to the network map.

### Node Feature Encoding

To infer the underlying thermodynamic properties and local flexibility, the feature vector *x*_*i*_ is engineered to capture sequence-encoded driving forces and structural proxies for molecular motions.

For all the residues at the interface, the node feature vector is 28-dimensional, of which the first 22 features define the local chemical identity, comprising a one-hot amino acid encoding, the Kyte-Doolittle hydrophobicity scale, and the formal charge. Quantifying the local hydrophobicity allows the network to identify the hydrophobic interactions that help anchor and stabilize the interface, as well as map the complementary binding pockets on structured partners. The formal charge captures the long-range electrostatic interactions that frequently drive the initial association and structural tethering of IDPs.

To account for the local flexibility, we use the B-factor, which is a well-established proxy for the thermal motion and fuzziness in the bound state. Additionally, the local backbone dihedral angles (*ψ, ϕ*) are calculated to allow the network to identify regions with transient secondary structural propensities that may facilitate binding.

Finally, each node is provided with spatial context through the minimum Euclidean distance to the nearest partner C*α* atom on the partner chain, encoding its proximity to the binding interface (e.g., a residue at 3 Å vs. 7.5 Å), along with an interface flag indicating the residue participates in the binding interface. Notably, each residue is assigned a binary chain identity flag (0/1) indicating its originating chain, which allows the model to distinguish residues belonging to different chains and correctly resolve pairwise interactions in multi-chain complexes. For the folded protein model, a 29^th^ dimension is initialized as an explicit node-level bias term.

### Message Passing

The model consists of a stack of three message passing layers with a hidden dimension of Each layer l takes the node embeddings ***h***^(*l*)^ and 3D coordinate positions ***h***^(*l*)^ as input, updating the embeddings to ***p***^(*l*+1)^:

#### Message Computation

First, for every edge (*j, i*) ∈ ***E***, a message ***m***_***ij***_ is computed that aggregates information between the receiving node *i* and the neighboring sender node *j*. Rather than relying on absolute spatial coordinates, the message function *ϕ*_*e*_ (a Multi-layer Perceptron (MLP)) is conditioned on the relative spatial vector between interacting residues, defined as **Δ*p***_***ij***_ = *p*_*i*_ − *p*_*j*_. By utilizing the relative position, the message passing operation ensures translation invariance. The message is formulated by concatenating sender features, receiver features, and their relative geometry:

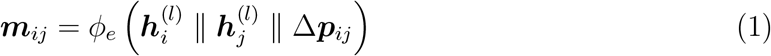

where ∥ denotes concatenation, and *ϕ*_*e*_ is a two-layer Multi-layer Perceptron utilizing a SiLU activation function, where 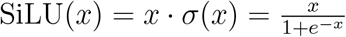

#### Neighborhood Aggregation

Once the pairwise messages are computed, each receiving node *i* gathers the structural and chemical information from its defined spatial neighborhood 𝒩 (***i***). To calculate this local environment, we employ a mean-pooling aggregation scheme:

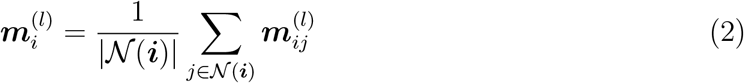

By averaging the incoming messages, the aggregated vector 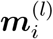, captures the overall physicochemical composition of the localized microenvironment, such as the general polarity, charge distribution, or hydrophobicity of a binding pocket.

#### Node Update

Following aggregation, each node *i* updates its latent representation by fusing its current state with the collective information from its neighborhood. The node embedding ***h***^(*l*)^ is concatenated with the aggregated message 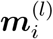 and passed through a node specific update function *ϕ*_*h*_ (an MLP with SiLU activation).

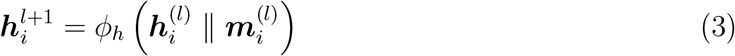

This ensures that the intrinsic identity is not lost in the neighborhood information. By concatenating the two vectors, the model allows the update function to weigh the residue’s individual properties against the collective chemical state of the surrounding interface, resulting in a refined embedding 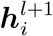 that represents the residue in its specific structural context.

#### Readout and Prediction

After L=3 message passing iterations, the final node embeddings 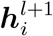 represent each residue as a product of both its intrinsic properties and its local environment. To generate a single prediction for the entire complex, a global mean pooling operation is applied as the readout function, aggregating all node-level embeddings into a fixed-size graph-level representation regardless of the number of interface residues:

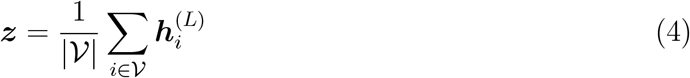

While this ensures the architecture can process complexes of varying sizes, the quality of the resulting embedding remains dependent on the distribution of graph sizes encountered during training. Complexes whose interface topology, in terms of number of nodes and edges, falls significantly outside the training distribution may not be reliably represented.

The global vector is then passed through a final prediction head consisting of a two-layer MLP with SiLU activation and a dropout layer to prevent overfitting. The output is a single scalar, representing the predicted binding energy of the protein-protein complex.

### Implementation Details

FlexENN was trained using an Adam optimizer with a learning rate of 1 x 10^−3^ for a batch size of 16 and 32 for the IDPs and the folded dataset, respectively, and 125 epochs. Model parameters were learned by minimizing the mean squared error (MSE) between the predicted and the experimental binding affinities across each batch. For clarity throughout the study, the model trained on the folded complexes is hereafter referred to as FlexENN-F, while the model trained on the IDP complexes is designated as FlexENN-I.

## Results and Discussion

### Limitations of the Ensemble Learning with Rosetta Features

The Protein Binding Energy Estimation (PBEE) framework ^31^ represents the latest attempt to combine Rosetta-derived energetic features with machine learning (ML). The algorithm employs a Super Learner (SL) architecture that stacks ten base models, including AdaBoost, decision trees, elastic net, extra trees, KNeighbors, linear regression, random forest, support vector regressor, and XGBoost, to leverage complementary inductive biases and enhance predictive robustness across diverse protein-protein complexes. Although the original study reported an RMSE of 1.98 kcal/mol and a Pearson correlation of 0.70 (Figure S4), the application of the framework to our benchmark set revealed significant limitations, as described next. A schematic overview of the SL training is provided in Figure S2. When applied to our benchmark set, the PBEE framework exhibited a systematic failure to discriminate between IDP complexes that share a common binding partner but differ substantially in the interface geometry (Table 4). For the *β*-catenin pairs, where 1JDH and 2GL7 differ by 3.0 kcal/mol in experimental binding affinity, PBEE predicted nearly identical values (−13.41 and −13.68 kcal/mol, respectively), failing to capture the energetics of the distinct interface configurations. PBEE also produced predictions that were more negative than the experimental affinities for IDP complexes, including MAP7 (predicted −14.2 kcal/mol versus −12.3 kcal/mol experimental) and the *β*-catenin complexes. Importantly, these failures were not confined to a single system but occurred across multiple benchmark complexes, indicating a systematic rather than an isolated issue. We therefore hypothesized that this behavior might arise from two critical factors in the training data: (1) the limited size of the overall dataset, which restricts the generalization across complex interaction landscapes, and (2) a severe representational bias against IDPs. With IDPs constituting only 4% (24 complexes) of the total dataset, the model likely failed to learn the distinct energetic principles governing disordered binding and is forced to extrapolate IDP energetics from a feature space dominated by folded proteins.

### Dataset Expansion Reveals Feature Collapse And Overfitting

To address the above limitations, we created two datasets, as described in the Methods section: one comprising 2,257 folded protein complexes (PBEE + Folded) and the other comprising 1,074 IDP complexes (PBEE + IDPs).

In principle, increasing the dataset diversity should improve generalization and mitigate the issues we were encountering. Even though the RMSE improved with both the folded and the IDP datasets, to 1.92 and 1.87 kcal/mol, respectively (Figure 6A, B), retraining PBEE on the expanded dataset resulted in mean-regression behavior, where predictions for binding free energies for all benchmark systems converged toward the mean values of the datasets (∼9.9 kcal/mol). Thus, while the model no longer produced the implausible values from above instead lost sensitivity altogether. From a practical point of view, such a model is unusable for comparative energetics analysis, as it cannot distinguish between different interfaces.

**Figure 6:**
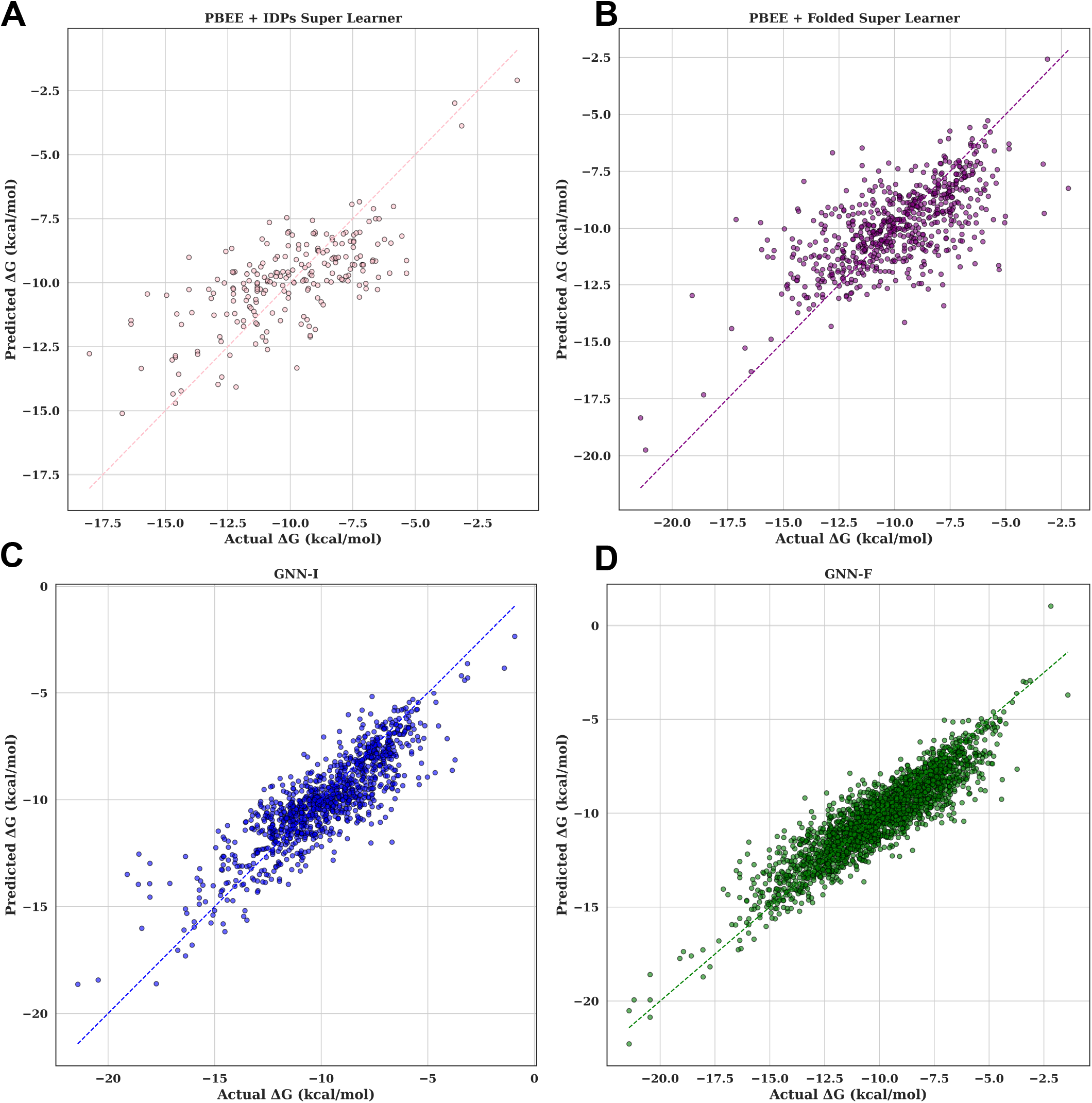
True versus predicted values generated from (A) the Super Learner trained on PBEE + IDPs dataset, (B) Super Learner trained on PBEE + folded dataset, (C) FlexENN-I trained on PBEE + IDPs dataset, and (D) FlexENN-F trained on PBEE + folded dataset.

We examined the learning behavior of the individual base models within the SL framework. In both the original PBEE study and in our retrained models, severe overfitting was present, especially for the Decision Trees, Extra Trees, and XGBoost, which consistently showed a training error of zero across all cross-validation folds while the validation errors plateaued, exhibiting a classic overfitting curve, shown in Figure S3.

To isolate whether this failure was due to the complex stacked architecture model or the input features themselves, we switched to individual ensemble learners, especially XGBoost, Decision Trees, and Extra Trees Regressor, using the same Rosetta energy descriptors. Despite using them independently, the overfitting persisted in the Decision Trees and Extra Trees Regressor. Even though XGBoost no longer showed a training error of zero, the predictions on our benchmark dataset were still at values near the mean ΔG of the dataset. For example, the model was unable to distinguish between the interdimer and lateral interfaces of the MT. The fact that distinct algorithms converge on similar failure modes strongly suggests that the limitation arises from the input representation rather than the learning architecture.

An additional and particularly concerning observation was the extreme sensitivity of Rosetta-feature-based models to minor changes in dataset composition. Removing as few as five randomly selected complexes from the training set resulted in dramatic degradation of prediction quality for the benchmark systems. Such instability indicates that learned decision boundaries are strongly influenced by individual data points, rather than reflecting robust, physically meaningful trends.

### Adaptive Graph Representations Overcome the Rigid Constraints of the Rosetta Features

The observed failure of Rosetta features to predict the IDP binding energies likely arises from a mismatch between rigid-body physical calculations and the thermodynamic reality of disordered binding. Traditional energy functions, such as Rosetta BETANOV16 (used in the PBEE method), were historically parametrized to balance high-resolution structural features against small-molecule thermodynamic data, thereby capturing proteins in a single, stabilized state. This approach is not well-suited for calculating IDP interactions because IDP binding often involves a shift within the broader conformational ensemble rather than a single native structure. Also, the strict distance-based scoring struggles to account for the fuzziness of interfaces where multiple interconverting conformations contribute to the total affinity.

We hypothesize that FlexENN handles the above limitations by functioning as an inherently adaptive model, evaluating the broader spatial context of the binding pocket rather than relying on a rigid set of deterministic constraints. Traditional physical scoring functions apply fixed, universal rules, where minor coordinate deviations, such as slight atomic clashes in a frozen crystal structure, often incur severe, unrealistic energy penalties. Our model bypasses this rigidity through three key architectural choices. First, by restricting the graph construction exclusively to the interface residues and modeling both their intra- and intermolecular edges, the network isolates the relevant microenvironment, capturing both the local structural integrity of the individual chains and the interactions between them. Second, by explicitly embedding dynamic structural descriptors (such as B-factors and secondary structure propensities) alongside sequence features, the network learns to contextually weigh static structural data against intrinsic disorder on a residue-by-residue basis. Finally, by utilizing relative distance vectors, rather than absolute spatial coordinates, the message-passing framework dynamically adapts to localized coordinate fluctuations, rendering the model more robust to the global conformational shifts that typically confound traditional algorithms.

### Predictive Performance of FlexENN

To evaluate our model’s capacity to predict protein-protein binding energies, we assessed its performance on our benchmark dataset comprising both folded and IDP complexes. We used the Root Mean Squared Error (RMSE) and the Pearson correlation coefficient (r) as our primary evaluation matrices (Figure 6, Table 1). To establish a robust comparative framework, we benchmarked FlexENN against several traditional Rosetta-feature-based baselines: the original PBEE model, PBEE models adapted to extended datasets (with more folded and IDP complexes), and traditional tree-based machine learning algorithms, including XGBoost, Decision Trees, and Extra Trees Regressor.

**Table 1:** Comparison of the Predictive Performance of our FlexENN Models with Other Methods.

| Model | RMSE (kcal/mol) | Pearson Correlation |
| --- | --- | --- |
| PBEE | 1.98 | 0.70 |
| PBEE + folded | 1.91 | 0.67 |
| PBEE + IDPs | 1.88 | 0.69 |
| XGBoost | 0.137 | 0.90 |
| Decision Trees | 0.00 | 1.00 |
| Extra Trees | 0.00 | 1.00 |
| FlexENN-F | 1.18 | 0.89 |
| FlexENN-I | 1.15 | 0.90 |

As an initial validation check, we monitored the training loss to prevent overfitting and dataset memorization. The tree-based models, XGBoost, Decision Trees, and Extra Trees failed this criterion, as their training loss converged to zero. A second validation check was implemented based on the folded complexes dataset to evaluate the model’s ability to differentiate the stronger tubulin interdimer interface (−14.9 ± 1.5 kcal/mol^50^) from the comparatively weaker lateral interface (−6.9 ± 0.4 kcal/mol ^50^). Our model (FlexENN-F) outperformed all baselines in this task (Table 2). While XGBoost and Decision Trees showed comparable differentiation, their severe overfitting rendered them unreliable for generalized predictive use.

**Table 2:** Comparison of the Predicted Binding Energy Values (kcal/mol) for the Tubulin Interdimer and Lateral Interfaces.

| System | True Values | FlexENN-F | PBEE + Folded | XGBoost | Decision Trees | Extra Trees |
| --- | --- | --- | --- | --- | --- | --- |
| Interdimer | $-14.9 \pm 1.5$ | -12.80 | -9.77 | -10.38 | -10.38 | -9.94 |
| Lateral | $-6.9 \pm 0.4$ | -5.88 | -8.94 | -7.31 | -8.63 | -9.31 |

### System True Values FlexENN-F PBEE + Folded XGBoost Decision Trees Extra Trees

The models trained on the folded dataset were subsequently tested on the CDC and NDC structures (the C and N terminal domains of DCX, respectively). Because the experimental ground truth for DCX includes energetic contributions from the entire protein, we averaged the models’ predictions for the two domains to ensure a fair comparison. In this test, the FlexENN-F model showed the most accurate predictions (predicted −10.37 kcal/mol against the experimental value of −10.8 kcal/mol) compared to the alternative base models, shown in Table 3.

**Table 3:** Comparison of the Predicted Binding Energy Values for DCX Across Different Models.

| System | True Value | FlexENN-F | PBEE + Folded | XGBoost | Decision Trees |
| --- | --- | --- | --- | --- | --- |
| CDC (6RF2) | DCX: $-10.8$ | $-8.84$ | $-9.73$ | $-8.82$ | $-10.43$ |
| NDC (6REV) | | $-11.9$ | $-9.51$ | $-9.02$ | $-10.65$ |
| **Avg. Value** | | $-10.37$ | $-9.62$ | $-8.92$ | $-10.54$ |

*Note: All values are reported in kcal/mol. DCX refers to the Doublecortin (NDC + CDC).*

**Table 4:** Comparison of the Predicted Binding Energy Values for IDP Complexes Across Different Models.

| System | True Values | FlexENN-I | PBEE | PBEE + IDPs |
| --- | --- | --- | --- | --- |
| Tau (6CVN) | −10.6 | −8.36 | −9.86 | −8.19 |
| MAP7 (7SGS) | −12.3 | −12.1 | −14.2 | −9.70 |
| PCNA-p21 (1AXC) | −9.7 | −9.42 | −9.07 | −9.39 |
| β-catenin-Tcf4 (1JDH) | −11.6 | −11.13 | −13.41 | −11.02 |
| β-catenin-Tcf4/BCL9 (2GL7) | −8.6 | −8.04 | −13.68 | −11.01 |

*Note: All values are reported in kcal/mol.*

Finally, because our initial work used the original PBEE to predict binding energies for the IDP complexes (tau and MAP7), we compared our IDP-specific model, FlexENN-I, against both the original PBEE and the extended PBEE variant, PBEE + IDPs (Table 4). It is important to note that, for tau, a direct comparison to the experimental values is precluded by a structural discrepancy: the benchmark experimental data reflects the full-length tau (with all four repeats), whereas our structural input (6CVN) represents only the isolated R2 repeat on MT. Therefore, we expected a deviation from the macroscopic experimental value in this case. For MAP7, FlexENN-I performed the best among all the tested models, yielding predictions closest to the experimental ground truth (predicted −12.1 kcal/mol compared to the experimental value of −12.3 kcal/mol).

We next assessed whether FlexENN-I generalizes beyond the MT system on three IDP complexes, 1AXC, 1JDH, and 2GL7, from distinct biological contexts. These complexes span more than 3 kcal/mol in experimental binding affinity and differ substantially in interface architecture, with 1AXC representing a short-motif bound to a homotrimeric clamp, and 1JDH and 2GL7 each comprising a long disordered region wrapping a folded scaffold.

On all three complexes, the FlexENN-I predictions fell within 0.6 kcal/mol of the experimental values. For 1AXC, the model predicted −9.4 kcal/mol compared to an experimental value of −9.7 kcal/mol. For 1JDH, the prediction was −11.1 kcal/mol compared to −11.6 kcal/mol. For 2GL7, the model predicted −8.0 kcal/mol compared to −8.6 kcal/mol (Table 4).

The pair of *β*-catenin complexes, 1JDH and 2GL7, provides a particularly informative test. Both share *β*-catenin as a binding partner, and both involve Tcf4 as one of the disordered partners, yet their binding energies differ by 3.0 kcal/mol, which is a consequence of the interface geometry. In 1JDH, Tcf4 alone engages with the *β*-catenin, whereas in 2GL7, both Tcf4 and BCL9 bind *β*-catenin independently at distinct sites, without contacting each other. FlexENN-I predicted a 3.1 kcal/mol gap between the two complexes, in close agreement with the 3.0 kcal/mol experimental difference. The model therefore distinguishes between the two *β*-catenin interactions based on the interface geometry rather than the binding partner identity, a discrimination capacity that is necessary for any binding energy model to be useful in practical applications.

### Limitations of the PDBbind Dataset

To assess the structural diversity captured by our dataset, we analyzed the distribution of graph sizes, quantified as the number of edges per complex, for both the PBEE + Folded and the PBEE + IDPs datasets. Across both datasets, the majority of samples are concentrated in the low-to-moderate range (below 300 edges). In both datasets, 97.5-97.6% of complexes contain fewer than 300 edges, and only 20 folded and 8 IDP complexes exceed 400 edges, corresponding to less than 1% of either dataset (Figures S5 and S6).

These results indicate that both datasets are dominated by complexes with small to moderate interfaces, whereas larger interfaces, such as those formed by multi-domain assemblies, or extended disordered regions wrapping around a folded partner, are under-represented. Importantly, this limitation is not specific to our model but reflects an inherent bias in datasets such as PDBbind, where large protein-protein interfaces are less frequently sampled than smaller complexes. Accordingly, predictions for complexes with higher edge counts may be unreliable, due to the scarcity of comparable examples during training. This highlights a broader challenge in data-driven modeling of protein-protein interactions, i.e., that the current datasets do not uniformly span the full range of biologically relevant interface sizes. Addressing this limitation will require the incorporation of more large-interface complexes or the development of strategies to better represent larger interfaces.

Notably, all benchmark complexes evaluated in this study, including the MT interdimer and lateral interfaces, the CDC and NDC for the folded systems, and the Tau-MT, MAP7-MT, PCNA-p21, *β*-catenin-Tcf4, and the *β*-catenin-Tcf4/BCL9 complexes, fall within the well-sampled range of their respective training distributions (as shown in Table 5 and Figures S5 and S6, ≤ 96th percentile in both cases), ensuring that the model evaluates structures within a range supported by sufficient training data.

**Table 5:** Number of Edges in the Graph Neural Network Representations of the Benchmark Proteins.

| System | Number of Edges |
| --- | --- |
| Interdimer | 272 |
| Lateral | 76 |
| CDC (6RF2) | 152 |
| NDC (6REV) | 2166 |
| Tau (6CVN) | 192 |
| MAP7 (7SGS) | 106 |
| PCNA-p21 (1AXC) | 220 |
| β-catenin-Tcf4 (1JDH) | 204 |
| β-catenin-Tcf4/BCL9 (2GL7) | 260 |

### Ablation Studies probe the robustness of the FlexENN architecture

To evaluate the efficiency and robustness of our proposed model architecture, we performed a series of ablation studies. By systematically removing individual node features, we constructed multiple model variants. This allowed us to quantify the impact of each node feature on overall performance and elucidate its specific contribution to the accurate prediction of binding energies of protein complexes.

We applied the previously established validation criterion to determine whether the model could accurately differentiate between the interdimer and lateral interfaces of MT (Figure S7). When we independently removed the node features, amino acid identity, hydrophobicity, charge, phi/psi angles, B-factor, and minimum distance between partner chains, the model failed to distinguish between the two interfaces. This failure highlights the model’s high sensitivity to information loss and demonstrates that none of the selected features are redundant. Instead, these results indicate that the distinction between the interdimer and lateral interfaces is not governed by a single dominant biochemical or structural property. Rather, their unique binding signatures emerge from a complex interplay of local geometry, electrostatics, and residue dynamics. FlexENN strictly requires the synergetic integration of all node features to construct a sufficiently rich structural representation capable of capturing the molecular landscape unique to each interface.

Next, we evaluated the feature reliance for the binding energy prediction of the globular domains of DCX (NDC and CDC; experimental baseline: −10.8 kcal/mol; FlexENN-F predicted value: −10.37 kcal/mol) (Figure S8). This analysis revealed a distinct hierarchy of importance for structured proteins. While removing the general physicochemical properties like charge (-9.68 kcal/mol), amino acid identity (−9.57 kcal/mol), and hydrophobicity (−9.35 kcal/mol) caused moderate predictive errors, structural constraints proved far more critical. Removing the psi angle or the minimum distance from the network changed the prediction to −9.02 and −8.9 kcal/mol, respectively. However, the most severe accuracy loss occurred in the absence of the phi angle and the B-factor, with values −8.36 kcal/mol and −7.86 kcal/mol, respectively, underscoring that the local geometry and the inherent residue flexibility govern the globular domain interactions.

We then shifted our focus to the intrinsically disordered MAP7 complex to evaluate the feature dependence of our IDP-specific FlexENN-I architecture (experimental value: −12.3 kcal/mol, and FlexENN-I predicted value: −12.1 kcal/mol) (Figure S9). Leaving out basic biochemical properties such as charge and hydrophobicity showed moderate deviations (−11.5 and −11.07 kcal/mol, respectively). However, the removal of the local conformational data caused severe predictive penalties: psi angle (−10.54 kcal/mol), phi angle (−9.76 kcal/mol), B-factor (−10.2 kcal/mol), and amino acid identity (−9.93 kcal/mol). The most drastic loss of performance occurred upon the removal of the minimum-distance feature (−7.72 kcal/mol), confirming that the precise spatial proximity constraints are the most critical determinants of the accurate modeling of highly flexible IDP-MT interactions.

Comparing the ablation profiles of the ordered DCX domains with those of the highly flexible MAP7 reveals a striking dichotomy in how the model perceives different binding mechanisms. For the intrinsically disordered MAP7, the network demonstrated an overwhelming dependence on the minimum distance feature. This aligns with the biophysical reality of IDPs, where energetic contributions are primarily driven by fuzzy, spatially distributed contacts. Conversely, the energetic profiling of the globular DCX domains was predominantly governed by the phi angle and the B-factor. Because structured proteins already possess a fixed overall shape, their binding strength is dictated by microscopic fine-tuning at the interface. They rely heavily on precise backbone angles and local residue flexibility to achieve a perfect structural fit. This stark contrast highlights a primary strength of our proposed model architecture, i.e., that, rather than applying a static uniform weighting to all inputs, the network dynamically prioritizes different geometric and physicochemical features based on the inherent structural nature of the complex. Therefore, the model successfully adapts to the distinct thermodynamic realities governing both ordered and disordered protein interactions.

## Conclusions

In this study, we successfully developed and validated an adaptive message passing graph neural network, FlexENN, capable of accurately predicting the binding energies of both the tubulin subunits in a MT and MT with MAPs. Crucially, our model demonstrated robust predictive power across a diverse structural spectrum, being able to adapt its feature prioritization to evaluate both rigidly folded domains and flexible IDPs.

A major contribution of this work is in providing a computational solution to address the failures of traditional, rigid-body scoring functions for disordered complexes. Rather than relying on strict physical parameters that artificially penalize natural coordinate variations, FlexENN evaluates the holistic spatial environment of the binding pocket. By incorporating structural descriptors directly into spatial representations, the model accurately accounts for interfacial dynamics using only static inputs. This allows the model to predict accurate binding energy values by appropriately weighing structural data against the local disorder.

While our model successfully infers conformational flexibility from structural descriptors, it does not explicitly break down the binding energy into its underlying enthalpic and entropic contributions. Importantly, this limitation is not unique to our approach: machine learning models, including the recent PBEE^31^ and SSIF-affinity^29^ approaches developed for folded proteins, similarly rely on structure-derived molecular descriptors and do not explicitly model these thermodynamic state functions. A key direction for future work is to move beyond single-conformation representations towards explicit modeling of the thermodynamic underpinnings of IDP binding, with the dual goal of improving the predictive accuracy and providing mechanistic insight into the driving forces governing the interactions responsible for the binding.

A related constraint stems from biases in the available training data, which do not fully capture the diversity of protein-protein interface architectures. While all benchmark systems considered in this work lie within the well-sampled regions of the training distribution, extrapolation to substantially larger or more complex interfaces may be less reliable. This limitation reflects a broader challenge associated with datasets such as PDBbind, rather than a model-specific issue, and highlights the need for expanded coverage or improved representations of structurally diverse complexes. Another natural extension of this work will be to incorporate trajectory data from molecular dynamics simulations, which provide time-resolved access to conformational ensembles and fluctuations. By integrating our adaptive message passing framework with larger, dynamic binding datasets, future iterations of this model could learn relationships between structural heterogeneity, entropy, and interaction energetics.

## Supporting information

Figures S1 to S9 supporting the main text analysis.

## Author Contributions

MI performed all the calculations and contributed to the writing of the manuscript. KOM carried out the experimental work to obtain the values. RID conceived the project, and KOM and RID contributed to the writing.

## Acknowledgement

This research was funded by the National Science Foundation (NSF) MCB-2527485. This work used the ACCESS program’s advanced computational resources through the allocation BIO210094 to RID.

## Data and Software Availability

All the data and input files with the code needed to reproduce this work are available on GitHub at https://github.com/DimaUClab/FlexENN.

## Supporting Information Available

Supporting Information: Figures of the second benchmark subset of IDPs, schematic of the PBEE SL setup, learning curves for XGBoost, Decision Trees, and Extra Trees, true versus predicted values for the SL trained on the original dataset, distribution of edge counts for PBEE + Folded and PBEE + IDPs datasets, Ablation results of FlexENN-F on interdimer and lateral interface, and DCX, Ablation results of FlexENN-I on MAP7.

## TOC Graphic

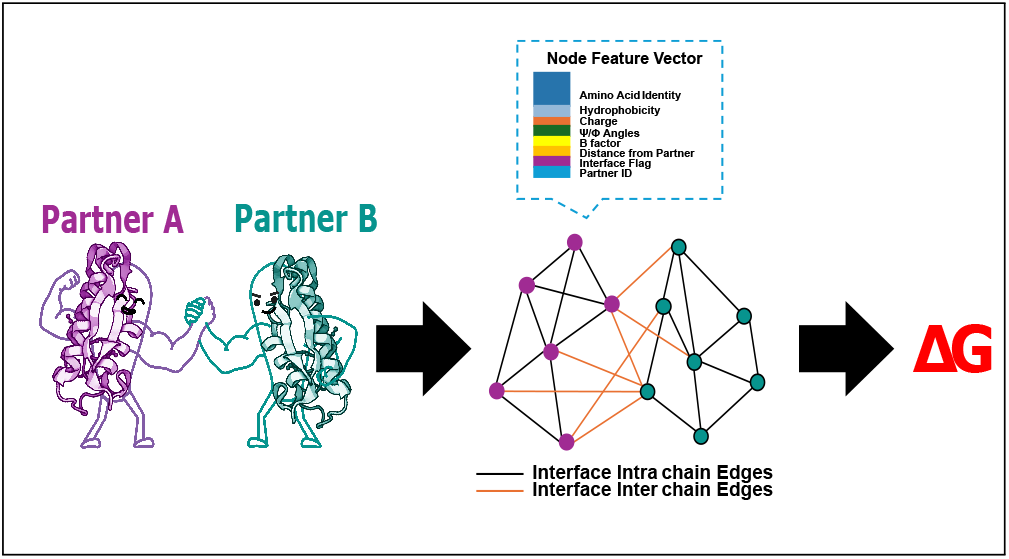

