## Supplementary material for "FlexENN: A Graph Neural Network for Binding Energy Prediction of Globular and Intrinsically Disordered Proteins": Figures S1 to S9 supporting the main text analysis.

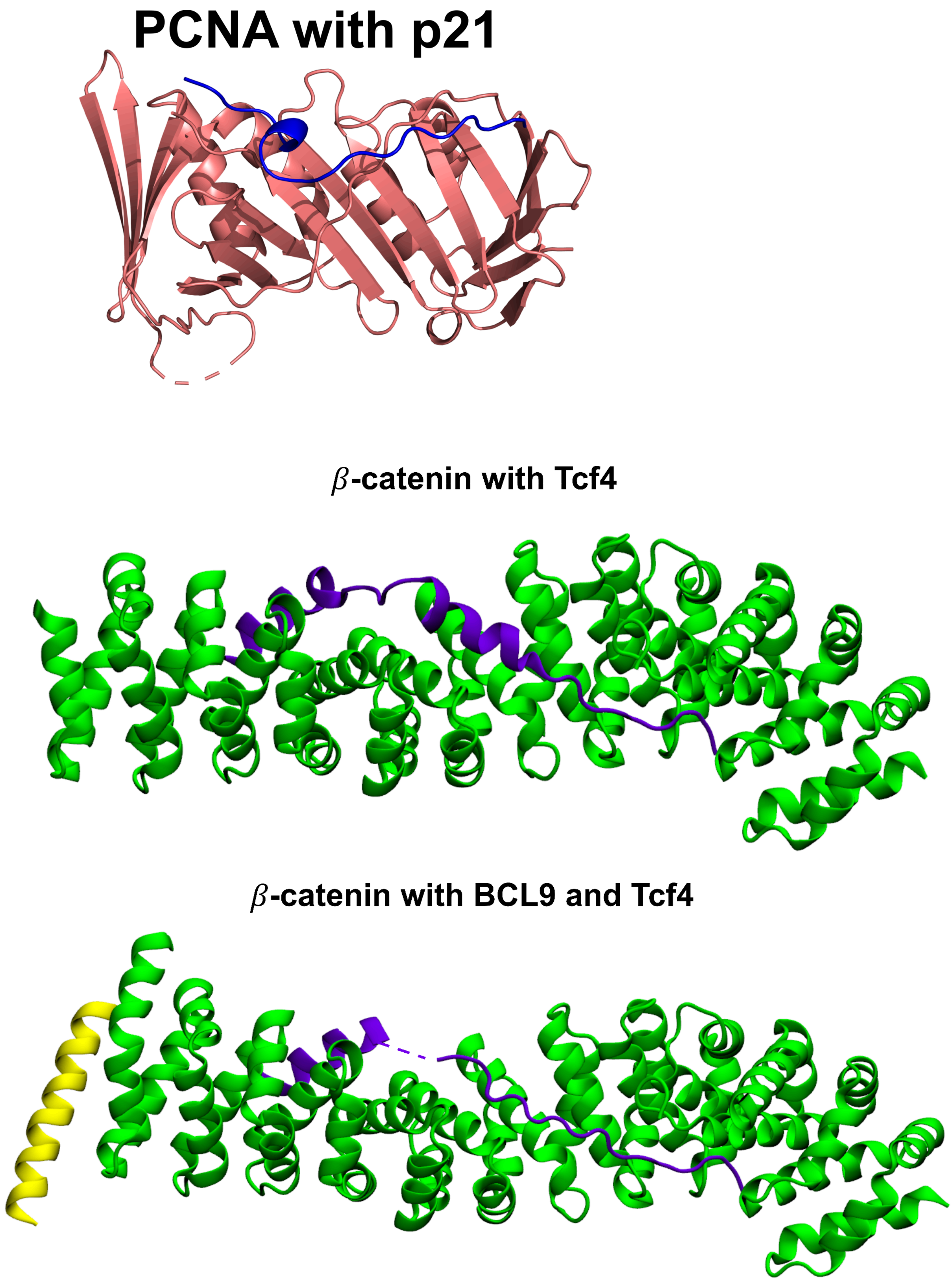


Figure S1. PDB structures of the second benchmark subset of IDPs.

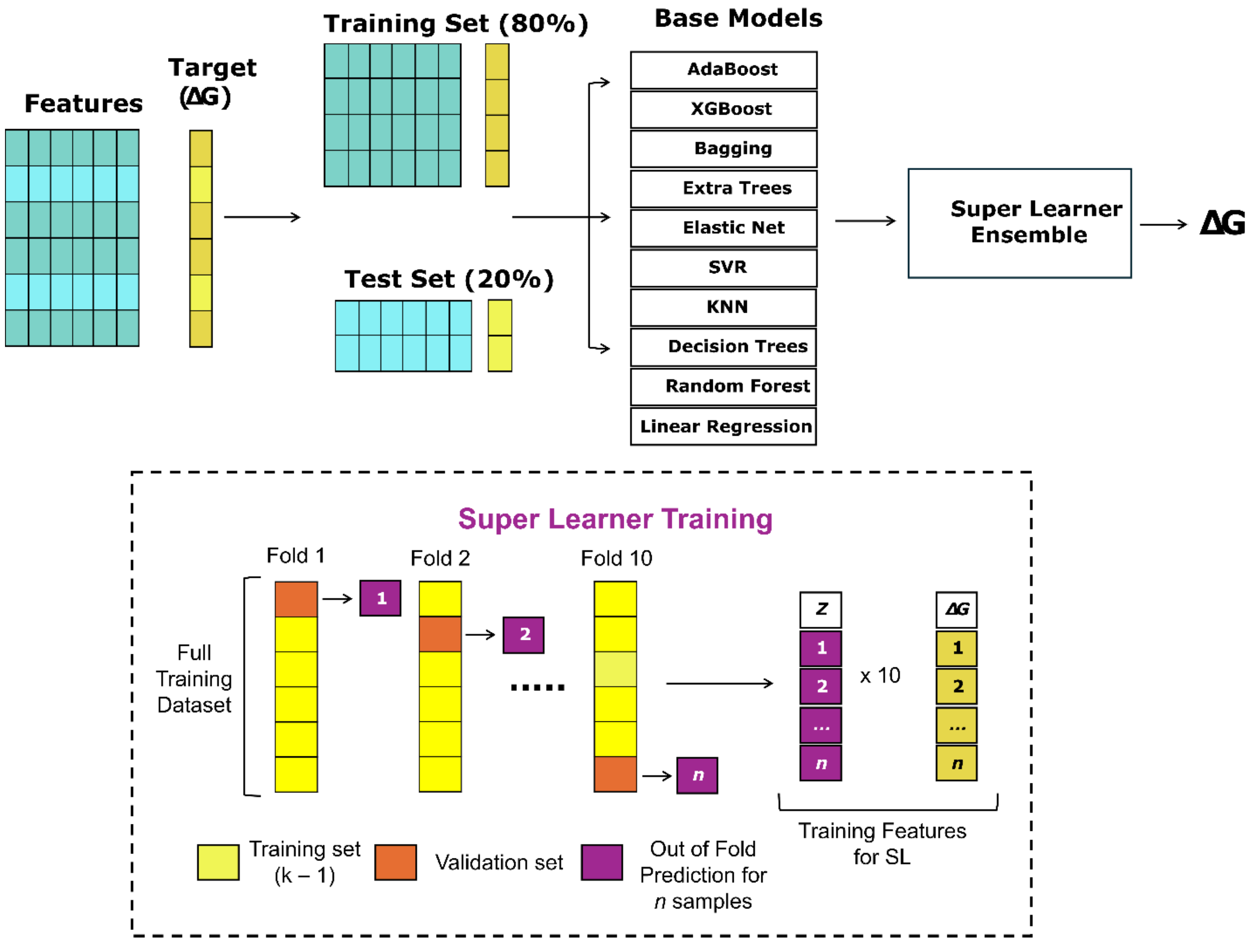


Figure S2. Schematic representation of the SL algorithm adapted from PBEE. The full training set is used to train 10 different ML algorithms. For the SL training, the dataset was split into 10-folds, and the subsets were trained on each ML algorithm and evaluated using k-fold cross-validation. All out-of-fold predictions were stored as a covariance matrix, and the model was fit to the subset and stored. The SL model was generated by combining predictions from each candidate using a weighted linear regression of their outputs on the full dataset.

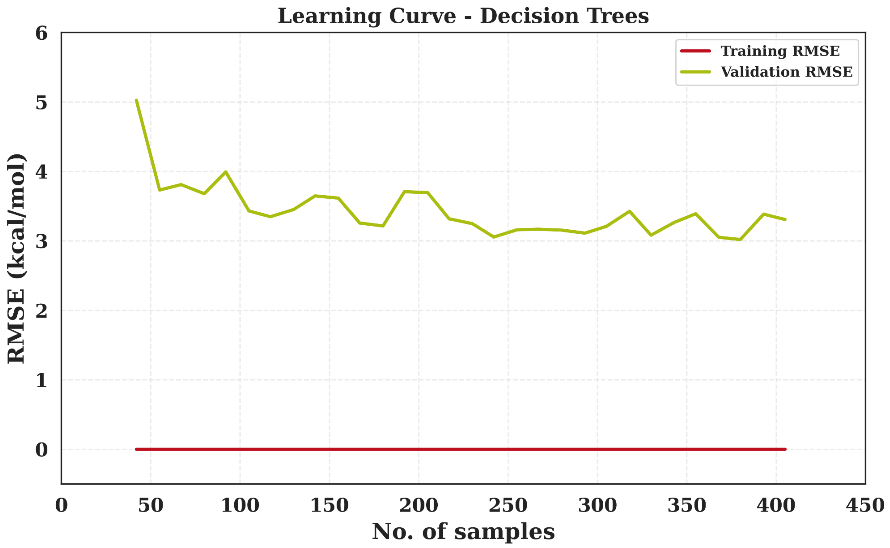
Learning Curve - Decision Trees
Training RMSE
Validation RMSE
RMSE (kcal/mol)
No. of samples
0
1
2
3
4
5
6
0
50
100
150
200
250
300
350
400
450

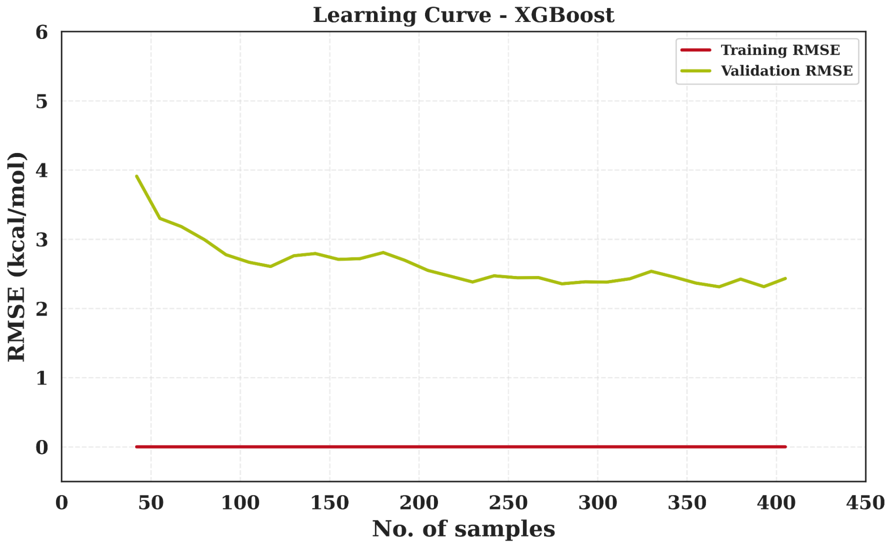
Learning Curve - XGBoost
Training RMSE
Validation RMSE
RMSE (kcal/mol)
No. of samples
0
1
2
3
4
5
6
0
50
100
150
200
250
300
350
400
450

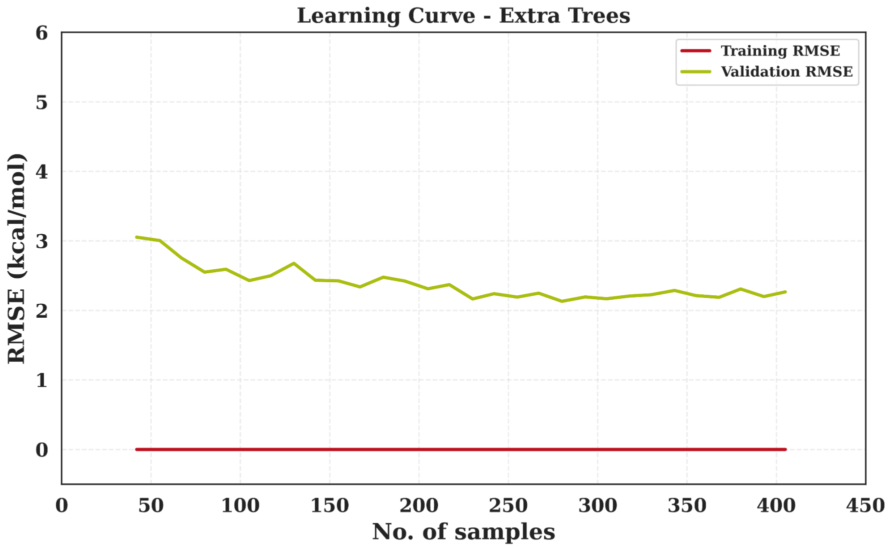


Figure S3. Learning curves for Decision Trees, XGBoost, and Extra Trees show clear overfitting, as evidenced by zero training error on the original PBEE Rosetta feature set.

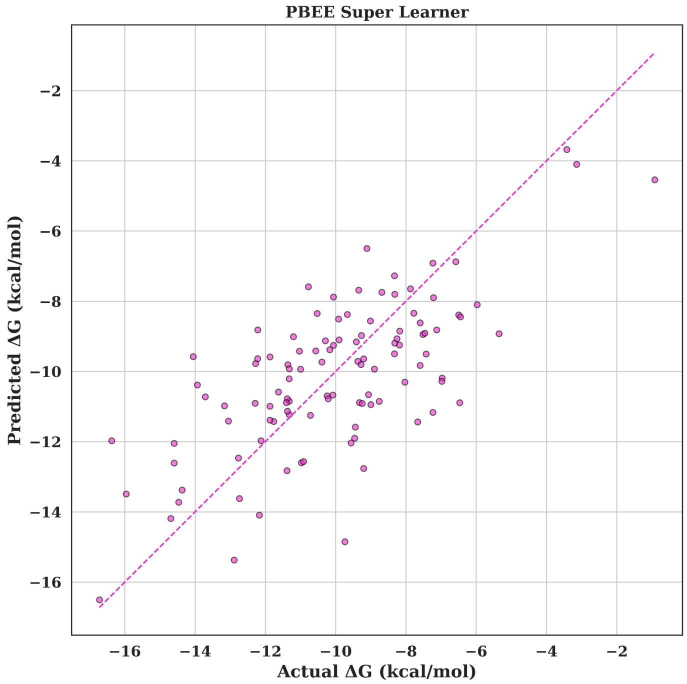


Figure S4. True versus predicted values generated from the Super Learner trained on the original PBEE dataset.

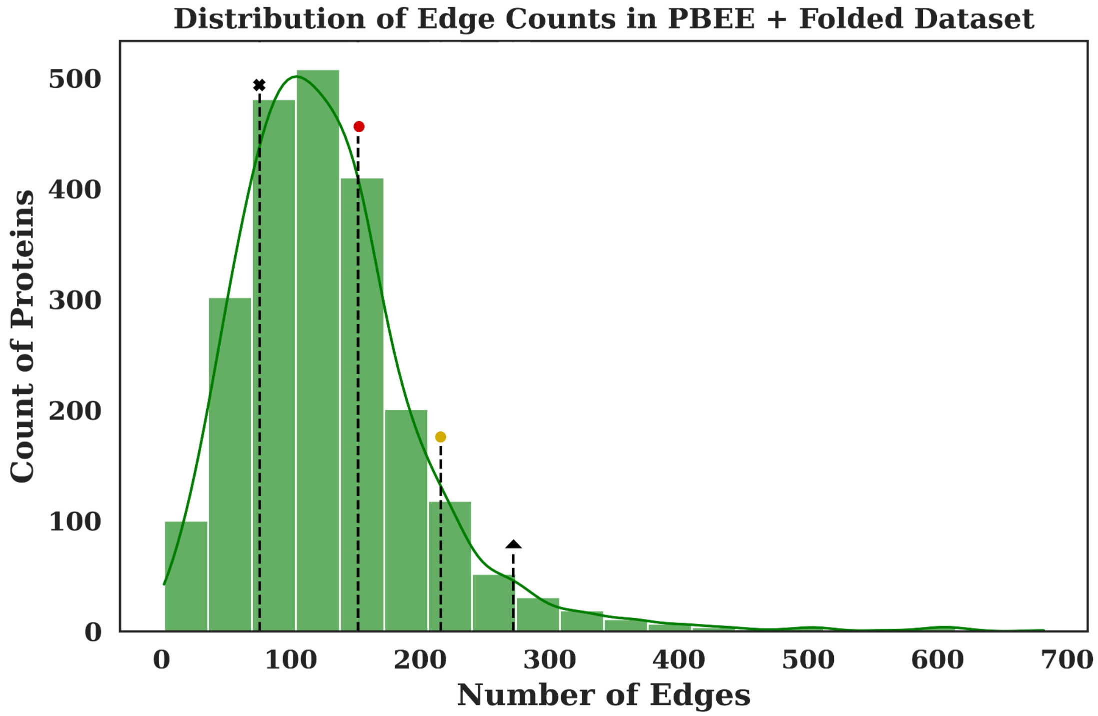


Figure S5. Distribution of edge counts per protein in the PBEE + Folded dataset. Vertical dashed lines indicate edge counts for reference benchmark folded MT-MAP structures: Tubulin lateral interface (✖, 76), CDC-MT (●, 152), NDC-MT (●, 216), and Tubulin interdimer/intradimer interface (▲, 272).

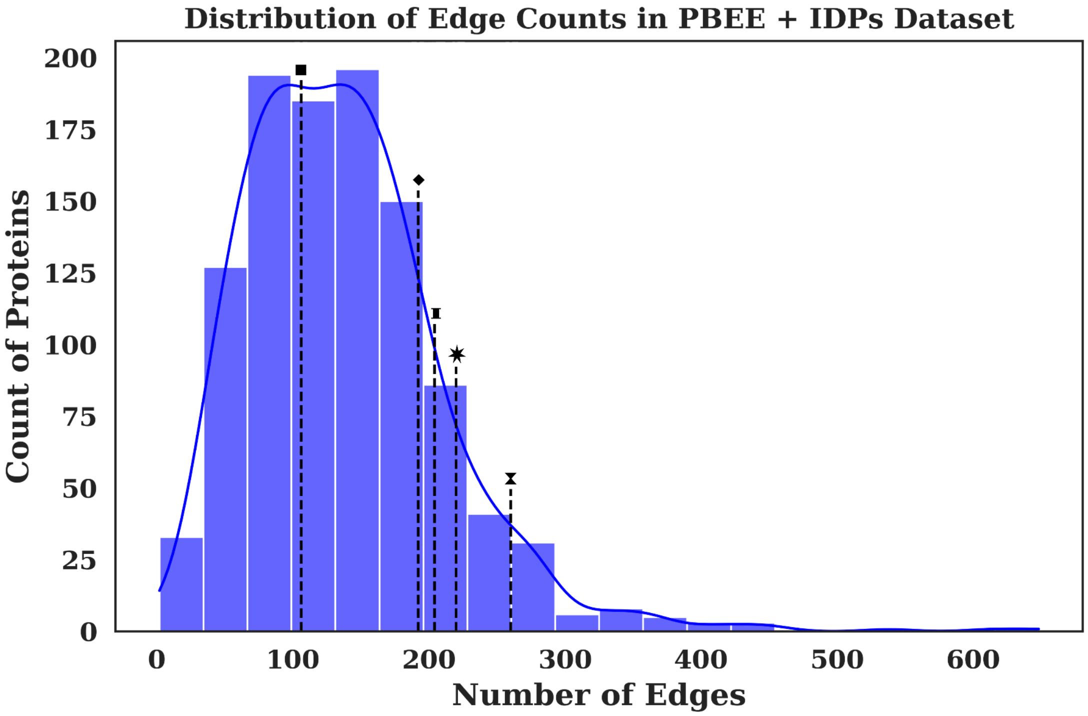


Figure S6. Distribution of edge counts per protein in the PBEE + IDPs dataset. Vertical dashed lines indicate edge counts for reference benchmark IDPs structures: MAP7-MT (■, 106), Tau-MT (◆, 192), $\beta$-catenin-Tcf4 (⌶, 204), PCNA-p21 (✶, 220), and $\beta$-catenin-Tcf4/BCL9 (⧗, 260)

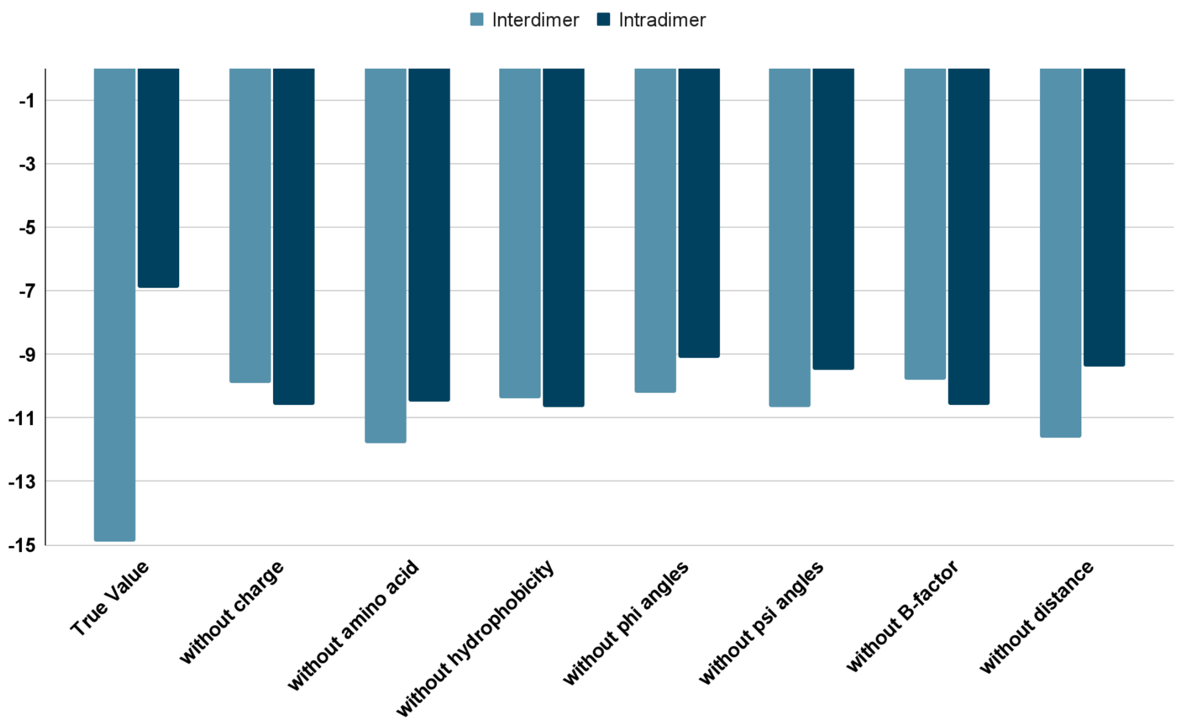


Figure S7. Ablation analysis of the FlexENN-F architecture for the prediction of ΔG values for MT's interdimer and lateral interface

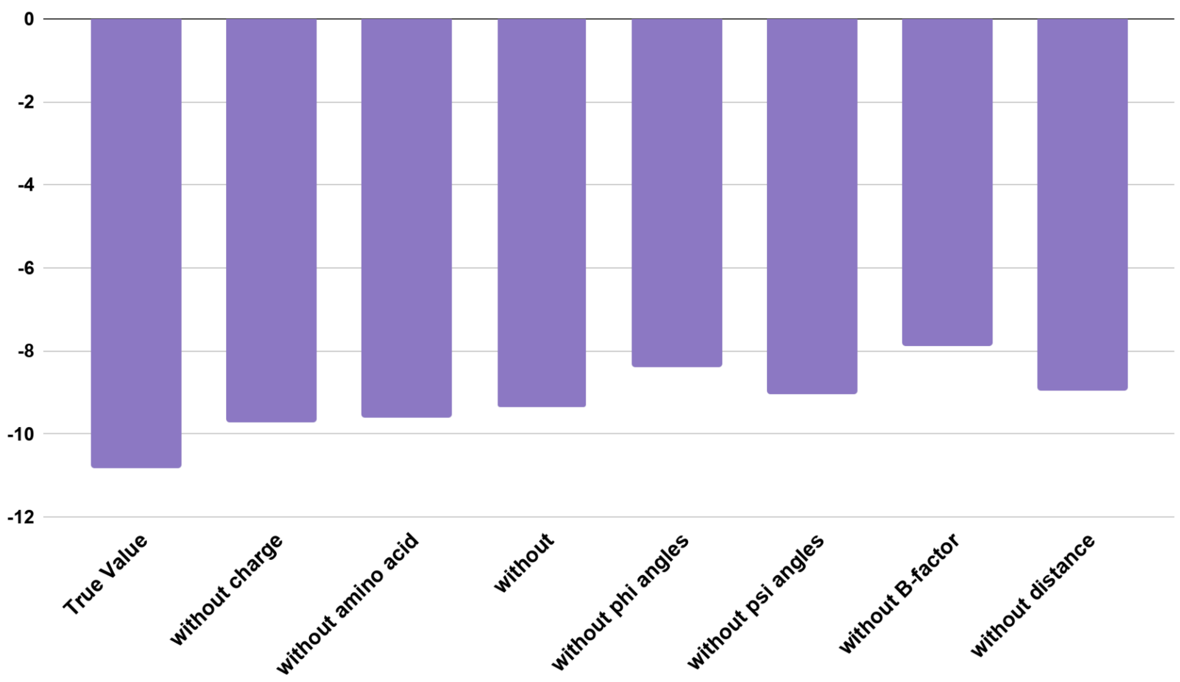

Figure S8. Ablation analysis of the FlexENN-F architecture for the prediction of ΔG values for DCX

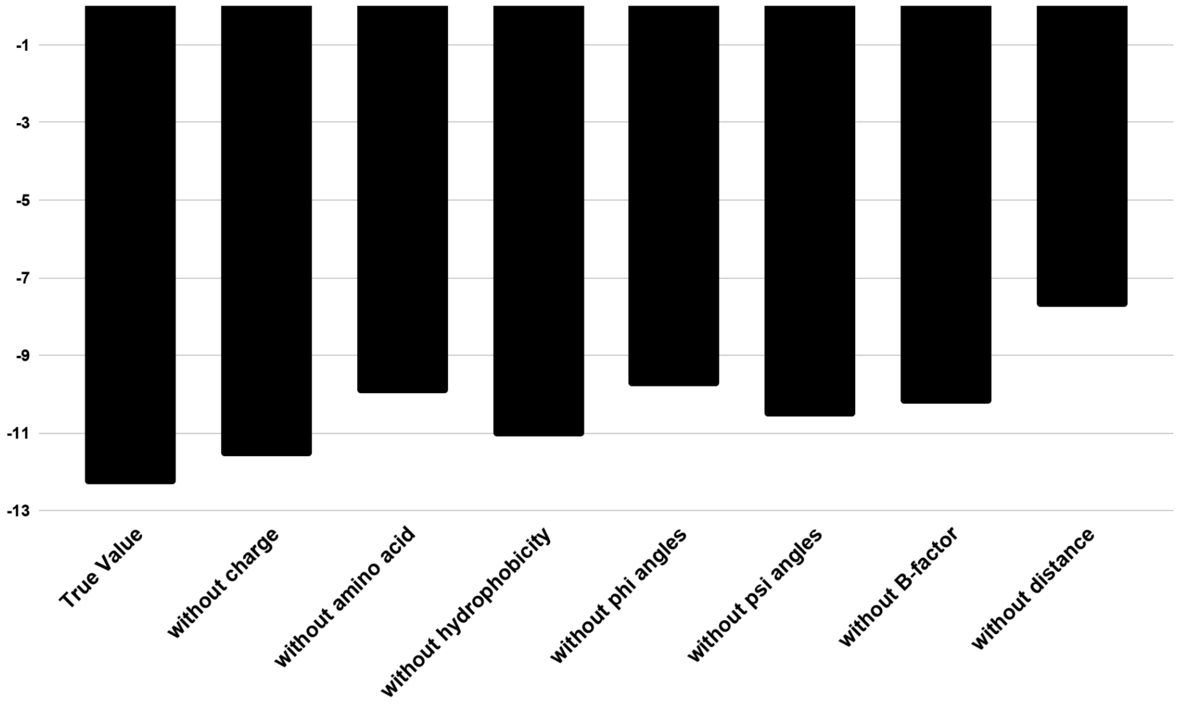


Figure S9. Ablation analysis of the FlexENN-I architecture for the prediction of ΔG values for MAP7
